# CAR-T Targeting of Mutant Calreticulin Establishes a Potentially Curative Stem Cell-Directed Therapy for Myeloproliferative Neoplasms

**DOI:** 10.64898/2026.02.09.704856

**Authors:** Daniel C Choi, Giovanni Medico, Irina V Lebedeva, Elisabeth K Nyakatura, Clarisse Kayembe, Franco Castillo Tokumori, Pouneh Kermani, Nassima Messali, Olivia Vergnolle, Abigail Taylor, Maria Mia Yabut, Urko del Castillo, Paul Balderes, Manuel Baca, Giorgio Inghirami, Joseph M Scandura

## Abstract

Myeloproliferative neoplasms (MPNs) are sustained by mutated hematopoietic stem cells (HSCs). Existing therapies fail to eliminate this compartment, leaving allogeneic HSC transplantation as the only curative option. Recurrent MPN driver mutations in calreticulin (*CALRmut*) generate a C-terminal neopeptide that requires cell-surface expression for oncogenic signaling, making it an attractive immunologic target. However, it remains unknown if CALRmut is uniformly displayed on all MPN HSCs within hematopoietic microenvironments. We generated huAB2, a high-affinity CALRmut-specific humanized antibody, to use as the targeting domain for chimeric antigen receptor (CAR)-T cells. We show that CALRmut is consistently displayed on functional MPN HSCs and accessible *in vivo*. huAB2 CAR-T cells eradicate MPN-propagating *CALRmut* HSCs in patient-derived tumor xenograft models without antigen escape while preserving coexisting normal human and host hematopoiesis. These findings establish CALRmut display as an obligate feature of MPN HSC fitness and support the feasibility of curative, non-transplant immunotherapy for *CALRmut* MPNs.

**Significance:** Therapies that eradicate cancer stem cells enable cure, but their feasibility is unknown. We establish an approach to potentially cure MPNs by proving mutant calreticulin to be a MPN stem cell marker that can be targeted by CAR-T cells to selectively wipe out disease in preclinical models of human MPNs.

## Introduction

Myeloproliferative neoplasms (MPNs) arise when a hematopoietic stem cell (HSC) acquires a driver mutation and gains a malignant fitness advantage over its normal counterparts. This leads to decades of progressive clonal dominance (1–3) that produces clinical disease and ultimately shortens survival (4). Because MPN HSCs are necessary and sufficient to initiate and sustain all MPN pathobiology (5,6), eliminating these cells is central to curing the disease. Most existing MPN therapies do not reliably target the malignant HSC compartment (7–9). As a result, most patients with MPNs lack options to alter the natural history of their disease.

An ideal immunologic target for durable and potentially curative disease modification must be selectively expressed by disease-propagating HSCs and indispensable for malignant fitness, to enable eradication of the malignant clone without immune escape and with selective sparing of normal hematopoiesis. Approximately thirty percent of patients with MPNs harbor driver mutations in calreticulin (*CALR*) (10) encoding a recurrent neoantigen (CALRmut) that uniquely satisfies these criteria. These mutations arise from deletions or insertions in exon 9 that uniformly generate a +1 frameshift, producing a highly conserved 44-amino acid C-terminal neopeptide that does not exist in normal cells (11). Wild-type CALR (CALRwt) is an endoplasmic reticulum (ER) resident chaperone that transiently binds nascent glycoproteins and is normally released during protein maturation via its native C-terminus (12,13). In contrast, CALRmut forms stable homodimers that associate with the thrombopoietin receptor MPL in the ER (14,15) but fail to dissociate with maturation due to the neo-C-terminus (16,17). The CALRmut/MPL complex traffics to the cell surface, where CALRmut functions as a pseudoligand for MPL, facilitating the assembly of second messengers that drive oncogenic JAK/STAT signaling (18–20). Importantly, surface expression of CALRmut is obligatory for malignant signaling (21), essentially linking immunologic accessibility to malignant fitness. Because MPL is expressed on HSCs (22), this biology predicts that CALRmut should be displayed on and essential to MPN HSCs, while completely absent from normal HSCs, thus positioning CALRmut as a compelling target for selective, disease-propagating HSC-directed immunotherapy.

Here, we engineered a fully humanized monoclonal antibody specific to surface CALRmut. We used its variable domain to generate CALRmut-targeted chimeric antigen receptor (CAR)-T cells active against CALRmut-expressing target cells *in vitro* and *in vivo*. We show that CALRmut-targeted CAR-T cells selectively eliminate primary patient-derived MPN HSCs while sparing normal hematopoiesis *in vivo*. These findings provide direct evidence that targeting CALRmut can eradicate disease-propagating HSCs and support the curative potential of this approach.

## Results

### Engineering huAB2, a humanized CALRmut-specific monoclonal antibody

Following immunization of mice with synthetic CALRmut C-terminal neopeptide, we generated a panel of 3,840 hybridomas and performed a tiered screen to identify antibodies with high affinity and specificity for CALRmut. Screening included binding to full-length CALRmut protein, the CALRmut C-terminal neopeptide, and CALRmut displayed in complex with MPL on the surface of cells. Counter-screens against cells lacking surface CALRmut were also conducted. From this pipeline, we identified AB2, a murine IgG2a antibody, as the top lead (Fig. 1A). AB2 was subsequently reformatted as a human IgG1 (huAB2).

**Figure 1:**
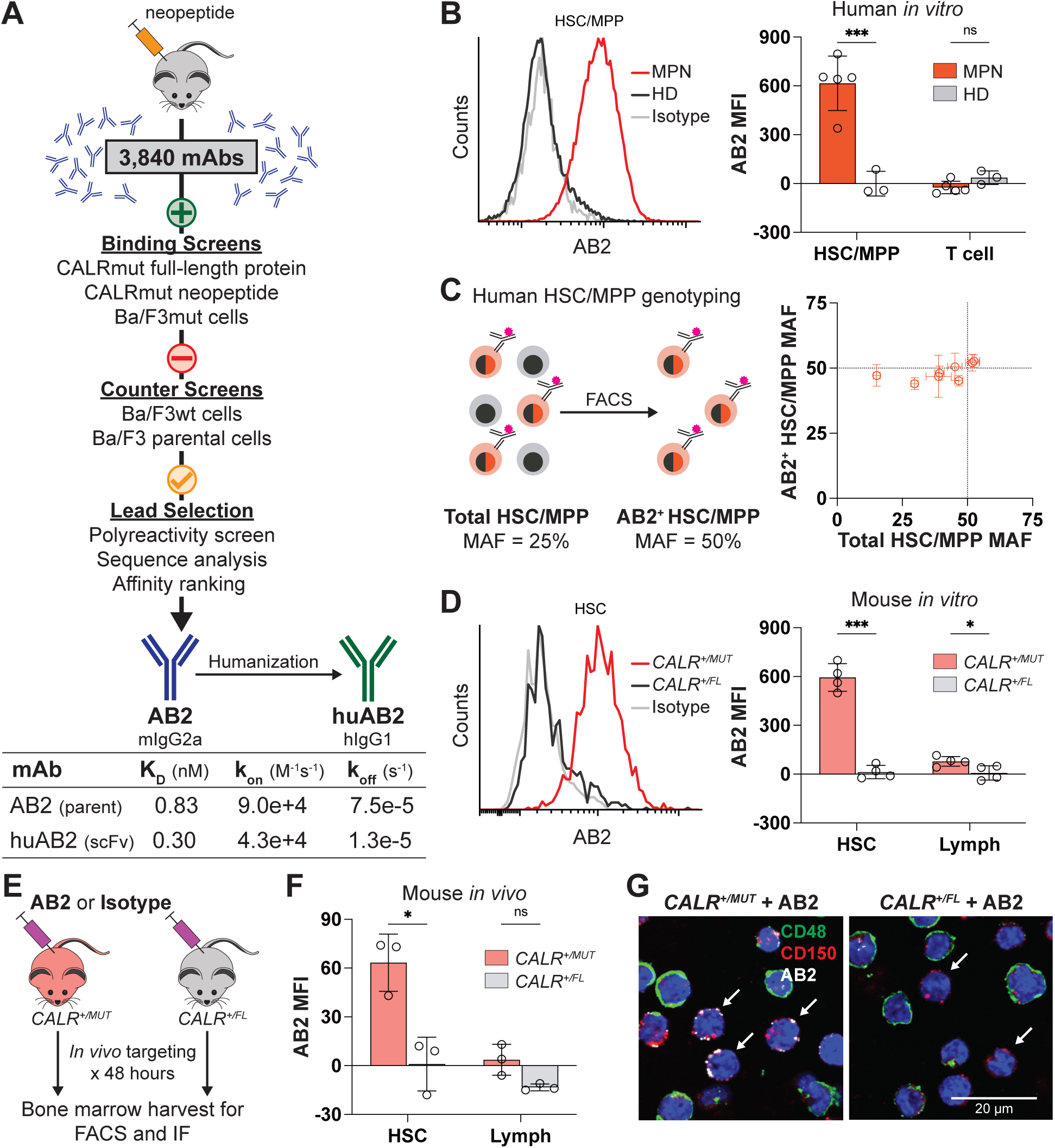
CALRmut-directed monoclonal antibody AB2 labels MPN HSCs *in vitro* and *in vivo*. **A)** Schematic representation of the screening approach taken to identify AB2. Rate constants for the AB2 parent murine antibody and the humanized AB2 (huAB2) scFv binding to synthetic CALRmut neopeptide are shown. **B)** Representative histograms and background-subtracted median fluorescence intensity (MFI) of *in vitro* AB2 staining of primary HSC/MPPs or T cells from MPN (n = 5) or healthy donor (HD, n = 3) peripheral blood. **C)** Enrichment of *CALR* mutation allele frequency (MAF) in sorted surface AB2-labeled HSC/MPPs (y-axis) compared to bulk HSC/MPPs (x-axis) from 7 MPN donors. MAF measurements from each individual donor are plotted with 95% confidence intervals (CI) indicated by error bars. **D)** Representative histograms and background (isotype)-subtracted MFI of *in vitro* AB2 staining of primary murine HSCs or lymphocytes from bone marrow of CALRmut-expressing *CALR^+/MUT^* mice (n = 4) or from *CALR^+/FL^*controls (n = 4). **E)** Experimental design for *in vivo* evaluation of AB2 targeting in *CALR^+/MUT^* mice. **F)** Background-subtracted MFI for *in vivo* AB2 staining of bone marrow HSCs or lymphocytes in *CALR^+/MUT^* mice (n = 3) relative to *CALR^+/FL^* controls (n = 3). **G)** Confocal immunofluorescence images of lineage-depleted bone marrow cells after *in vivo* AB2 staining. Shown is the *in vivo* AB2 staining pattern for murine HSCs (Lin^-^CD48^-^CD150^+^, white arrowheads) from *CALR^+/MUT^* (left) and *CALR^+/FL^* controls (right). Abbreviations: Monoclonal antibody (mAb), immunofluorescence (IF). Aggregate MFI data are presented as means with error bars representing standard deviation (SD) and each point representing one donor (*in vitro*) or mouse (*in vivo*). * p < 0.05, *** p < 0.001, ns = not significant (p ≥ 0.05) (t-test).

Using biolayer interferometry, we determined that AB2 binds the CALRmut C-terminal neopeptide with sub-nanomolar affinity both as the parent murine antibody and as a humanized single-chain variable fragment (scFv), with favorable on- and off-rates (Fig. 1A). To assess binding to CALRmut displayed in its physiologic context with MPL at the cell surface, we engineered Ba/F3 parental cells to co-express human MPL with either human CALRmut (Ba/F3mut) or CALRwt (Ba/F3wt). Fluorescence-activated cell sorting (FACS) analysis demonstrated robust binding of AB2 to Ba/F3mut cells, without detectable binding to Ba/F3wt cells (Supplementary Fig. S1A,B). These data confirm that AB2 selectively recognizes CALRmut presented with MPL at the cell surface.

### AB2 selectively recognizes surface CALRmut on primary patient-derived MPN cells

We next evaluated AB2 binding *in vitro* to CD34^+^ hematopoietic stem and progenitor cell (HSPC) subpopulations via FACS using blood samples collected from patients with *CALRmut* MPNs or from healthy donors (HD). AB2 selectively bound CD45^dim^CD34^+^CD38^-^CD45RA^-^ HSCs and multipotent progenitors (HSC/MPPs) from MPN samples, but not HSC/MPPs from HD samples, nor MPL^-^CD3^+^ T cells from the same donors (Fig. 1B and Supplementary Fig. S1C). Importantly, AB2 selectively labeled the most immature immunophenotypic HSCs (CD45^dim^CD34^+^CD38^-^CD45RA^-^CD90^+^) (23) representing putative disease-propagating HSCs in MPN samples (Supplementary Fig. S1D). Thus, AB2 recognizes surface-expressed CALRmut on MPN HSC/MPPs.

*CALR* driver mutations in MPNs are almost always heterozygous (10,24), such that a sample comprised exclusively of *CALRmut* cells is expected to have a mutation allele frequency (MAF) of 50%. We found that the majority of the MPN HSPC samples we analyzed had *CALR* MAFs of ∼50%, consistent with *CALRmut* predominance (Supplementary Table S1). To determine whether AB2 selectively labels *CALRmut* HSPCs, we generated defined mixtures of HSPCs from MPN and HD samples, producing populations with *CALR* MAFs ranging from 15%–53% (mean 40%). From these mixed samples, AB2^+^ HSC/MPPs were isolated by FACS and *CALR* MAF was quantified by droplet digital PCR (ddPCR) in both the total and AB2^+^ HSC/MPP fractions (Fig. 1C). Across all mixtures, the AB2^+^ HSC/MPP fraction was consistently enriched to a mean *CALR* MAF of 49% (±4%), approaching the theoretical maximum for a homogenous population of heterozygous mutated cells. Therefore, AB2-marked HSC/MPPs carry the *CALRmut* genotype.

### AB2 labels surface CALRmut on primary MPN HSCs *in vivo*

Although *CALR* is broadly expressed *in vivo*, surface display of CALRmut is restricted to rare cells that co-express MPL, which include HSCs and select hematopoietic progenitor populations (25). We therefore sought to determine whether AB2 can selectively label CALRmut-expressing HSCs not only *in vitro* but also *in vivo*, within their native bone marrow niche. To enable *in vivo* tracking, we generated heterozygous *CALRmut* knock-in mice (*CALR^+/MUT^*), in which a single allele of murine *Calr* bearing the human CALRmut C-terminal neopeptide sequence is expressed from the endogenous murine *Calr* locus following *Vav1*-Cre mediated recombination (26). *In vitro*, AB2 selectively bound murine HSCs (Lin^-^cKit^+^Sca1^+^CD48^-^CD150^+^) (27) isolated from the bone marrow of *CALR^+/MUT^* mice, but not lymphoid populations from the same mice or hematopoietic cells from *CALR^+/FL^*control mice (Fig. 1D, Supplementary Fig. S1E).

To assess targeting *in vivo*, we infused fluorescently conjugated AB2 into *CALR^+/MUT^*or control *CALR^+/FL^* mice and analyzed hematopoietic tissues 48 hours later (Fig.1E). At this time point, AB2 administration had no measurable effect on peripheral blood counts (Supplementary Fig. S1F). FACS analysis demonstrated selective AB2 labeling of bone marrow HSCs in *CALR^+/MUT^* mice, with no detectable labeling above background of HSCs in *CALR^+/FL^* controls or in lymphoid populations (Fig. 1F, Supplementary Fig. S1G). Direct visualization by confocal immunofluorescence microscopy confirmed AB2 labeling of HSCs from *CALR^+/MUT^*bone marrow (Fig. 1G). AB2 labeling was also observed in CD41^hi^ megakaryocyte-biased HSCs (Mk-HSC) (28) and in Lin^-^cKit^+^Sca1^-^ myeloid progenitor cell (HPC) populations, but not in peripheral blood lineages such as T cells, B cells, or platelets (Supplementary Fig. S1H). These findings show that AB2 selectively binds surface-expressed CALRmut on MPN HSCs both *in vitro* and *in vivo*, supporting its potential as a targeting platform for MPN HSC-directed immunotherapy.

### Functional CALRmut-targeted CAR-T cells were generated from huAB2

To generate a CALRmut-targeted CAR, we cloned the heavy- and light-chain variable domains of huAB2 into a scFv containing an N-terminal hemagglutinin (HA)-tag. This targeting domain was fused to a 4-1BB costimulatory domain and the CD3ζ signaling domain (Fig. 2A). We introduced this CAR construct into primary human T cells via a lentiviral vector. Across multiple T cell donors, the mean vector copy number was 1.1 (range 0.9–1.2) and mean transduction efficiency, as reported by HA-tagged CAR expression, was 65% (range 56%–72%) (Supplementary Fig. S2A). The resultant CAR-T cells were expanded several hundred-fold using anti-CD3/anti-CD28 antibody beads (Supplementary Fig. S2B).

**Figure 2:**
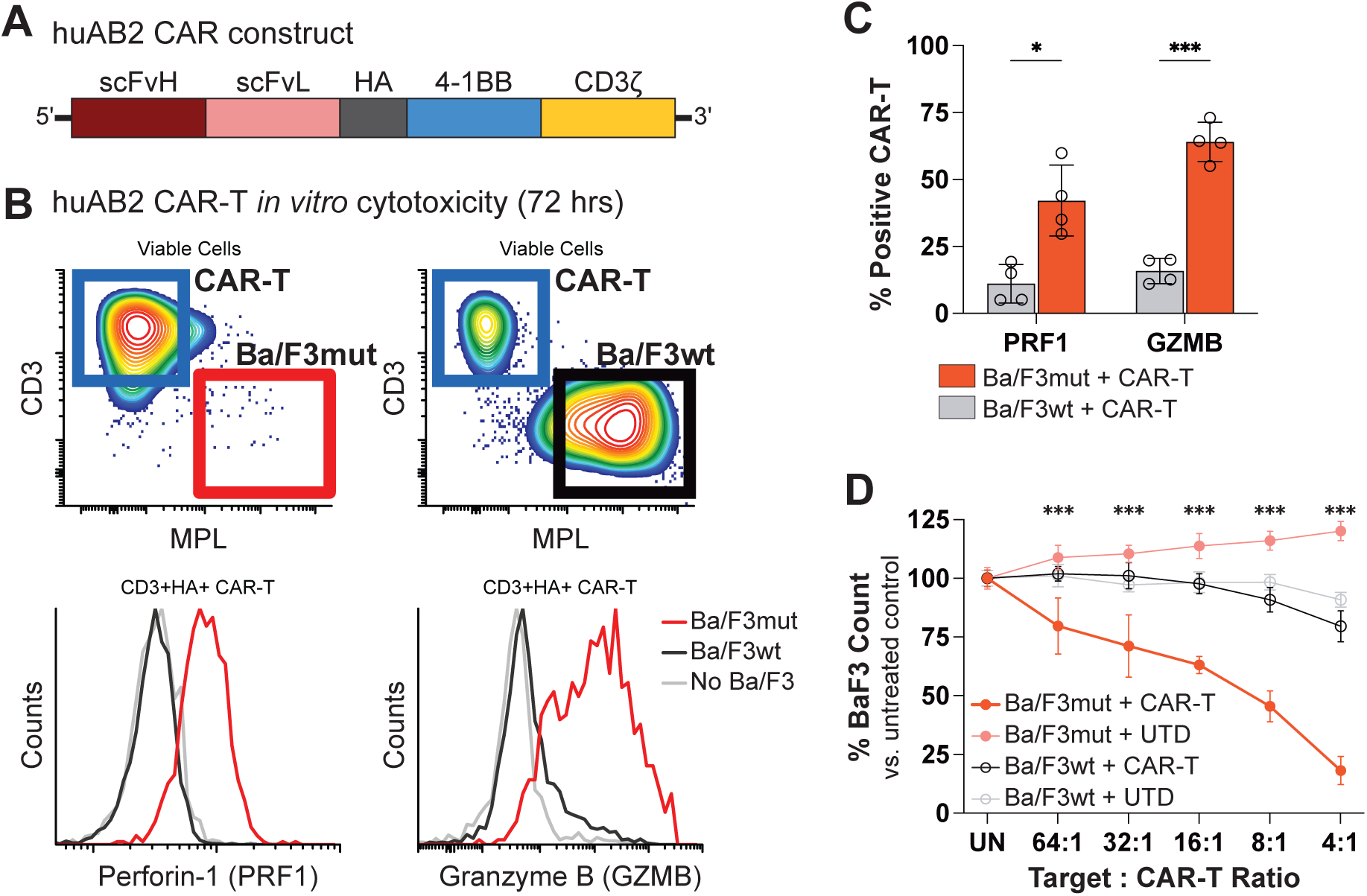
huAB2 CAR-T cells selectively kill CALRmut target cells *in vitro*. **A)** Schematic of the huAB2 CAR construct delivered via lentiviral transduction to generate CAR-T cells. **B)** Flow cytometry gating strategy used to identify huAB2 CAR-T cells and Ba/F3mut or Ba/F3wt cells in co-culture. Representative histograms show intracellular perforin-1 (PRF1) and granzyme B (GZMB) expression in CAR-T cells following 72 hours co-culture with Ba/F3mut targets or Ba/F3wt controls. **C)** Quantification of huAB2 CAR-T cells expressing PRF1 or GZMB after 72 hours co-culture with Ba/F3mut or Ba/F3wt cells, with 4 technical replicates per condition. **D)** Effector-to-target-dependent cytotoxicity of huAB2 CAR-T cells or untransduced T cell controls (UTD) against Ba/F3mut (mut) or Ba/F3wt (wt) cells over 72 hours *in vitro*, normalized untreated control culture cell counts. Abbreviations: scFvH (scFv heavy chain), scFvL (scFv light chain). Mean cell counts from triplicate co-cultures at each effector-to-target ratio are shown with statistically significant differences between Ba/F3mut and Ba/F3wt counts after CAR-T treatment noted. * p < 0.05, *** p < 0.001, non-significant comparisons not shown (t-test).

To assess antigen-specific activation, huAB2 CAR-T cells were co-cultured with Ba/F3mut or Ba/F3wt cells. The cytotoxic markers perforin-1 and granzyme B were robustly expressed by the majority of huAB2 CAR-T cells upon exposure to Ba/F3mut cells, whereas minimal expression was observed with Ba/F3wt exposure, confirming CALRmut antigen-specific activation (Fig. 2B,C). Consistent with this target-induced activation, huAB2 CAR-T cells mediated dose-dependent killing *in vitro* of Ba/F3mut cells but not Ba/F3wt cells (Fig. 2D). Untransduced T cell (UTD) controls showed no cytotoxic activity against either Ba/F3 cell line, confirming CAR-mediated killing and the absence of cross-species toxicity. Thus, huAB2 CAR-T cells can be efficiently generated, are specifically activated by surface CALRmut, and mediate potent and selective cytotoxicity *in vitro*.

### huAB2 CAR-T cells eradicate CALRmut target cells and prolong survival *in vivo*

To assess the efficacy of huAB2 CAR-T cells against CALRmut-expressing target cells *in vivo*, we developed a CALRmut-driven murine tumor model by implanting Ba/F3mut cells into immunodeficient NOD.Cg-*Prkdc^scid^Il2rg^tm1Wjl^*Tg(CMV-IL3,CSF2,KITLG)1Eav/MloySzJ (NSGS) mice (Fig. 3A) (29). In this aggressive model, Ba/F3mut cells rapidly expand *in vivo* (Supplementary Fig. S3A), infiltrating the blood, bone marrow, and spleen, and uniformly leading to death within 3 weeks (Supplementary Fig. S3B).

**Figure 3:**
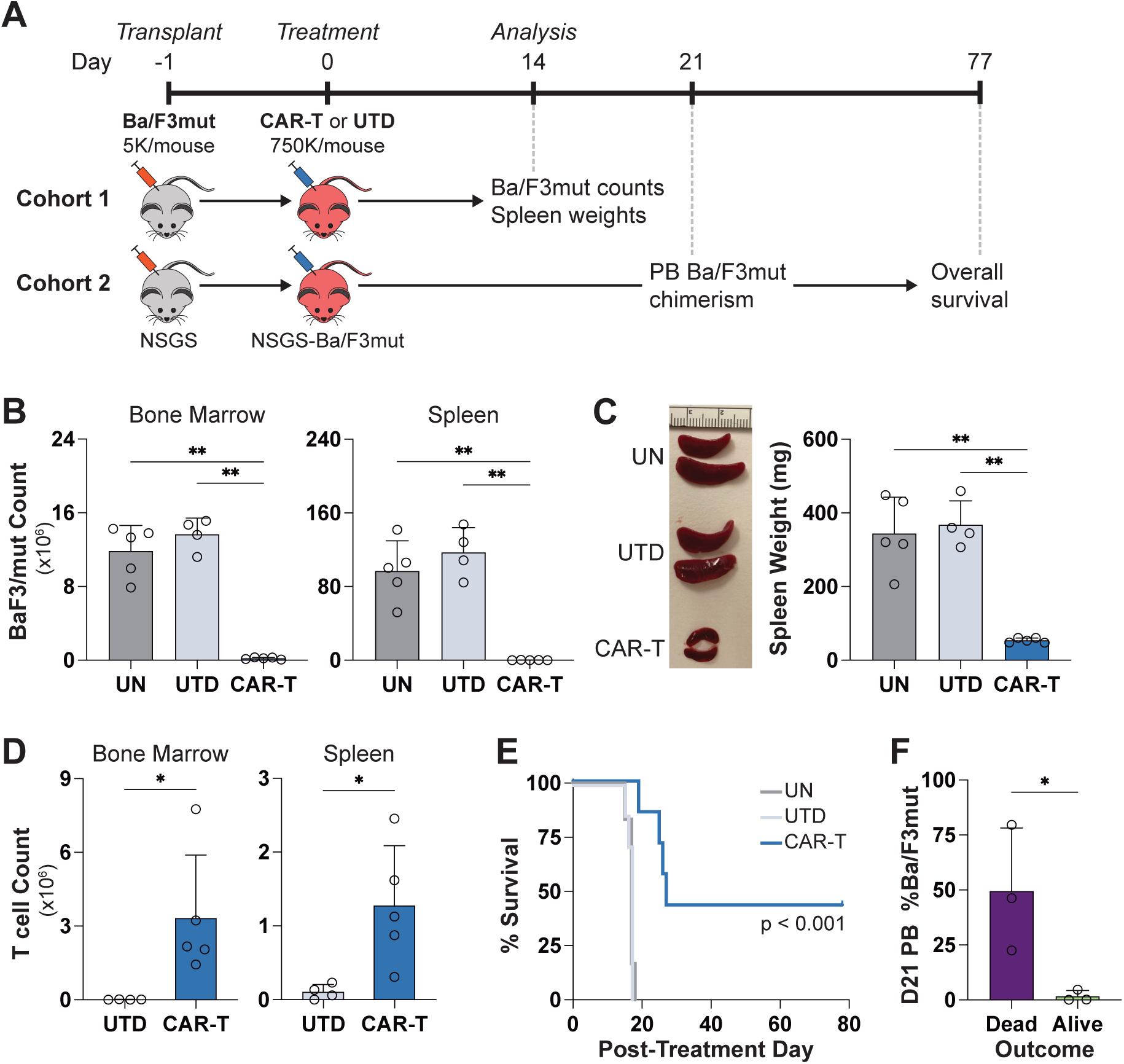
huAB2 CAR-T cells kill surface CALRmut-expressing Ba/F3mut cells *in vivo*. **A)** Experimental design for *in vivo* evaluation of huAB2 CAR-T cells versus UTD controls in the NSGS-Ba/F3mut murine tumor model. **B)** Mean engrafted Ba/F3mut cell counts in bone marrow and spleen of NSGS-Ba/F3mut mice treated with CAR-T cells (n = 5) or UTD (n = 4), or left untreated (UN, n = 5). **C)** Representative spleens (left) and mean spleen weights (right) from UN mice and those treated with CAR-T cells or UTD. **D)** Mean human T cell counts in bone marrow and spleen of NSGS-Ba/F3mut mice treated with CAR-T cells or UTD. **E)** Kaplan-Meier survival curves for NSGS-Ba/F3mut mice in UN (n = 6), UTD-treated (n = 7), and CAR-T-treated (n = 7) groups, with statistical significance assessed by log-rank test. **F)** Mean Ba/F3mut chimerism in peripheral blood (PB) of six CAR-T-treated NSGS-Ba/F3mut mice from the survival cohort at post-treatment day 21. Mice are stratified by survival outcome at analysis on post-treatment day 77. Error bars represent SD and each point represents an individual mouse. * p < 0.05, ** p < 0.01, non-significant comparisons not shown (t-test).

Ba/F3mut recipient mice were treated with huAB2 CAR-T or UTD controls 24 hours after tumor cell implantation. To analyze the effects of huAB2 CAR-T cells on tumor burden, an initial cohort of mice was euthanized at post-treatment day 14, prior to the demise of control mice receiving UTD or no T cells (UN). Treatment with huAB2 CAR-T cells significantly depleted Ba/F3mut cells in the bone marrow, spleen, and peripheral blood of the recipient mice compared to UTD-treated controls (Fig. 3B, Supplementary Fig. S3C). Spleen size was also significantly reduced in the CAR-T-treated mice compared to controls (Fig. 3C). huAB2 CAR-T cells expanded and persisted in the bone marrow and spleen of recipient mice whereas UTD controls were largely absent at analysis (Fig. 3D). The persistence of huAB2 CAR-T cells at post-treatment day 14 raised the possibility of long-term anti-CALRmut activity in recipient mice.

A second cohort of NSGS mice was transplanted with Ba/F3mut cells as described above to evaluate the impact of huAB2 CAR-T cells on survival (Fig. 3A). Treatment with huAB2 CAR-T cells significantly prolonged the survival of recipient mice compared to controls receiving UTD or UN (median survival 27 days for huAB2 CAR-T vs. 17 days for UTD and UN, p < 0.001, Fig. 3E). All mice in the UTD and UN control groups died by post-treatment day 18, and one mouse in the CAR-T group died on post-treatment day 20. We analyzed Ba/F3mut burden in peripheral blood collected from the remaining 6 CAR-T-treated survivors on post-treatment day 21. Results were dichotomous with significant Ba/F3mut burden detected in 3 mice (suggesting treatment failure) and virtually undetectable Ba/F3mut burden in the other 3 mice (Fig. 3F). All 3 mice with significant Ba/F3mut burden on day 21 died within one week, whereas the other 3 mice with minimal Ba/F3mut burden survived until scheduled euthanasia almost 3 months post-treatment. Therefore, huAB2 CAR-T cells mediate potent cytotoxic activity against CALRmut-expressing cells *in vivo* and can even prolong survival in a highly aggressive disease model.

### huAB2 CAR-T cells selectively target MPN HSPCs *in vitro*

Physiologic surface expression of CALRmut on primary MPN HSPCs is substantially lower than the enforced expression in the Ba/F3mut model. To evaluate huAB2 CAR-T cell activity against primary MPN cells, we first assessed *in vitro* killing of HSPCs isolated from 7 patients with *CALRmut* MPNs (Supplementary Table S1) and 7 HD controls. Treatment with huAB2 CAR-T cells *in vitro* selectively depleted HSPCs from MPN samples with minimal effect on HSPCs from HD samples compared to UTD treatment (Supplementary Fig. S4A).

HSPCs from MPN samples can include normal cells with those harboring *CALRmut*, as evidenced by HSPC MAFs <50% from select donors (Supplementary Table S1). To confirm preferential targeting of CALRmut-expressing HSPCs within these samples, we quantified *CALR* alleles by ddPCR following exposure to huAB2 CAR-T cells. Treatment with huAB2 CAR-T cells selectively depleted *CALRmut* alleles relative to *CALRwt* alleles, whereas both alleles were uniformly preserved upon UTD treatment (Supplementary Fig. S4B). This indicates that huAB2 CAR-T cells can selectively eliminate MPN HSPCs with physiologic surface CALRmut expression.

### huAB2 CAR-T cells eliminate MPN HSPCs, including immunophenotypic HSCs, *in vivo*

To evaluate huAB2 CAR-T cell efficacy *in vivo* against primary MPN HSPCs, we established a patient-derived tumor xenograft (MPN PDTX) model using HSPCs from 4 individuals with *CALRmut* MPNs (Supplementary Table S1). Robust MPN HSC engraftment has been historically challenging in xenograft systems (30–32), necessitating specialized graft or recipient modifications (33,34). To enable the engraftment of unmodified primary MPN HSCs, we developed an immune-deficient recipient with reduced endogenous HSC competition by backcrossing NSGS mice into the NOD.Cg-*Kit^W-41J^Tyr^+^Prkdc^scid^Il2rg^tm1Wjl^*/ThomJ (NBSGW) (35) background (NSGS-cKit^W-41J^). The donor MPN HSPC grafts had *CALR* MAFs ranging from 43%–51%, consistent with near-complete derivation from heterozygous CALRmut clones. MPN HSPCs rapidly engrafted recipient mice with a mean human chimerism of 83% (±20%) observed in the bone marrow within 14 days of implantation (Supplementary Fig. S4C).

Three days after implantation, MPN PDTX mice were treated with huAB2 CAR-T cells or UTD controls and euthanized on post-treatment day 14 for tissue analysis (Fig. 4A,B). Treatment with huAB2 CAR-T cells significantly depleted MPN donor-derived human CD33^+^ myeloid cells and HSPC subpopulations in the bone marrow of recipient mice relative to UTD treatment (Fig. 4C, Supplementary Fig. S4D). Critically, huAB2 CAR-T treatment markedly reduced MPN donor-derived immunophenotypic HSC/MPP populations that include disease-propagating MPN HSCs (Fig. 4D). Consistent with these findings, *CALRmut* alleles were virtually eliminated from bone marrow HSC/MPPs of CAR-T-treated mice yet preserved in UTD-treated controls (Supplementary Fig. S4E). Histopathologic analysis of the PDTX bone marrow confirmed the eradication of CALRmut-expressing cells in the CAR-T-treated mice relative to UTD controls (Fig. 4E).

**Figure 4:**
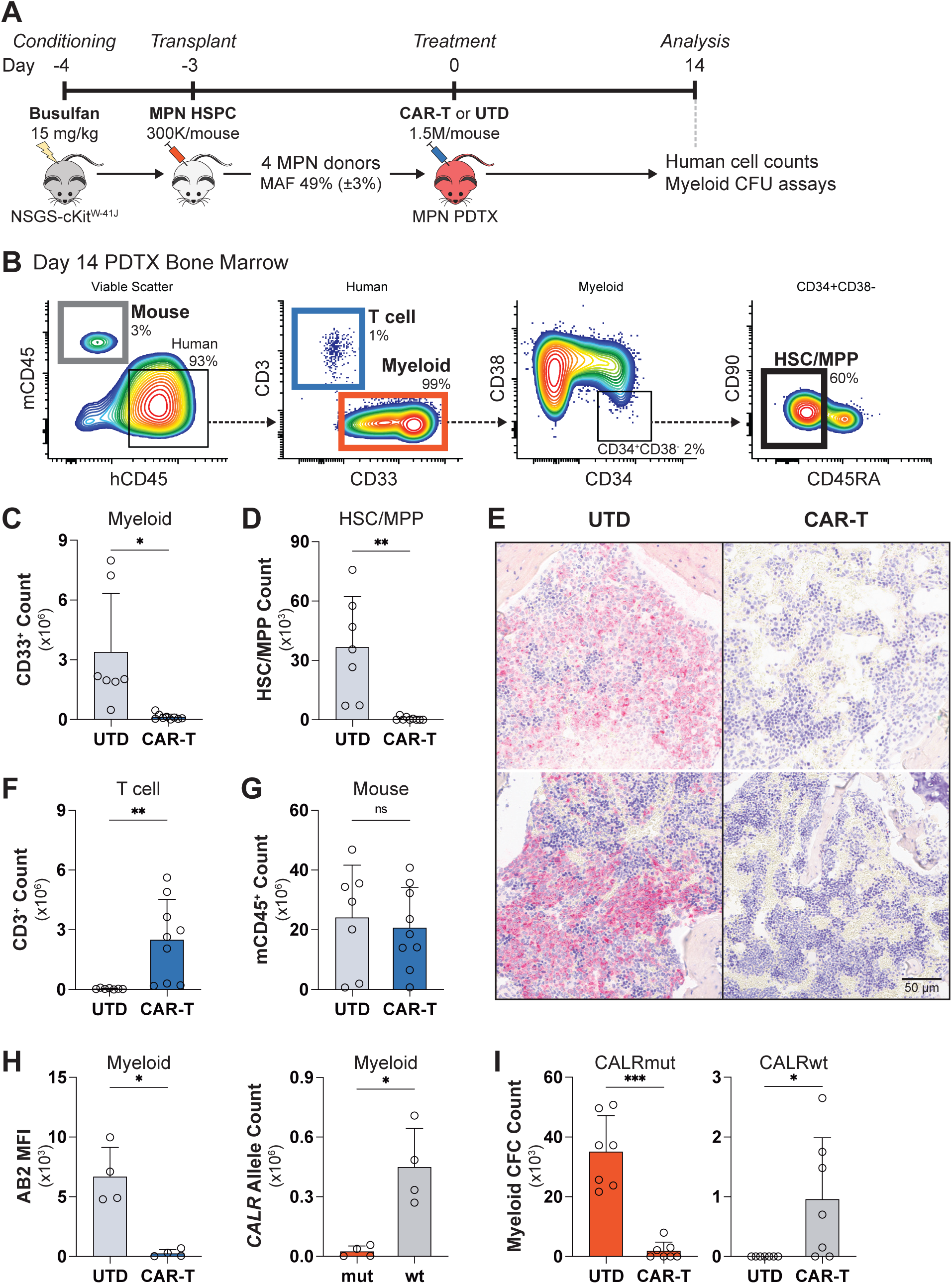
huAB2 CAR-T cells eradicate primary human MPN cells *in vivo*. **A)** Experimental design for *in vivo* evaluation of huAB2 CAR-T cells against primary MPN HSPCs in MPN patient-derived tumor xenograft (MPN PDTX) models. **B)** Flow cytometry gating strategy used to identify recipient-derived mouse cells, treatment-derived T cells, and MPN donor-derived myeloid cells and HSC/MPPs in bone marrow of MPN PDTX mice. Murine and human CD45 are designated mCD45 and hCD45, respectively. **C)** Mean human CD33^+^ myeloid cell counts in bone marrow of MPN PDTX mice 14 days after treatment with CAR-T cells (n = 9) or UTD (n = 7). **D)** Mean human HSC/MPP counts in MPN PDTX bone marrow 14 days after treatment. **E)** Representative immunohistochemistry of post-treatment MPN PDTX bone marrow showing absence of CALRmut-expressing cells (in red, via AB2 staining) in two CAR-T-treated mice compared to two UTD-treated controls. **F)** Mean human T cell counts in MPN PDTX bone marrow 14 days after treatment. **G)** Mean recipient-derived murine leukocyte counts MPN PDTX bone marrow 14 days after treatment. **H)** Left: AB2 MFI showing level of surface CALRmut expression on residual human myeloid cells present in MPN PDTX bone marrow after CAR-T or UTD treatment. Right: Mean *CALRmut* (mut) and *CALRwt* (wt) allele counts determined by ddPCR in residual human myeloid cells sorted from CAR-T-treated MPN PDTX bone marrow. **I)** Mean number of myeloid colony forming cells (CFCs) grown *in vitro* from bone marrow of CAR-T- or UTD-treated MPN PDTX mice, stratified by *CALRmut* and *CALRwt* genotypes. Error bars represent SD and each point represents an individual mouse. * p < 0.05, ** p < 0.01, *** p < 0.001, ns = not significant (p ≥ 0.05) (t-test).

huAB2 CAR-T cells expanded and persisted in the bone marrow of treated PDTX mice until post-treatment day 14, while UTD-T cells were largely absent at this time point (Fig. 4F). Recipient-derived murine leukocyte counts in the bone marrow were comparable between the CAR-T- and UTD-treated groups (Fig. 4G), indicating that huAB2 CAR-T cells killed MPN targets without detectable off-target toxicity to host hematopoiesis. Similar patterns of MPN cell depletion, CAR-T cell persistence, and recipient cell sparing were seen in the spleen, demonstrating robust and selective treatment effects across hematopoietic tissues (Supplementary Fig. S4F).

### Residual normal HSPCs are spared by huAB2 CAR-T cells in MPN PDTX mice

Despite profound depletion of *CALRmut* cells and alleles, low-level human cell engraftment remained detectable in all huAB2 CAR-T-treated MPN PDTX mice. To determine whether these residual cells represented antigen-escape MPN clones or spared normal hematopoiesis, we directly assessed surface CALRmut expression by AB2 staining and *CALR* genotype by ddPCR. Residual donor-derived cells lacked detectable AB2 staining and were almost exclusively *CALRwt* (Fig. 4H). Together, these data exclude antigen-loss immune escape by MPN clones and indicate that the residual human cells are donor-derived normal cells untargeted by huAB2 CAR-T treatment.

To functionally assess the hematopoietic capacity and donor clonal origin following treatment, we performed myeloid progenitor colony-forming cell (CFC) assays on bone marrow harvested from MPN PDTX mice and genotyped individual colonies. In UTD-treated MPN PDTX mice, donor-derived CFCs were uniformly *CALRmut*, consistent with persistence of MPN hematopoiesis and the absence of detectable normal progenitors (Fig. 4I). In contrast, CAR-T-treated MPN PDTX mice showed a marked reduction in total donor-derived CFCs, reflecting profound depletion of MPN hematopoiesis in these animals. Notably, *CALRwt* CFCs were recovered in 5 of 7 CAR-T-treated MPN PDTX mice evaluated. These findings indicate that huAB2 CAR-T treatment selectively ablates *CALRmut* progenitors, enabling the functional re-emergence of residual normal hematopoietic clones that were below the limit of detection in untreated disease.

### huAB2 CAR-T cells preserve healthy hematopoiesis *in vivo*

To directly test whether huAB2 CAR-T cells selectively target MPN hematopoiesis while sparing normal hematopoiesis *in vivo*, we generated chimeric PDTX mice by implanting mixed donor grafts comprised of equal numbers of HSPCs from male patients with *CALRmut* MPN and female HD HSPCs into NSGS-cKit^W-41J^ recipients. Chimeric PDTX mice were treated with huAB2 CAR-T cells or UTD controls as previously described and analyzed 14 days later (Fig. 5A). MPN donor-derived cells were quantified from FACS-sorted subpopulations by ddPCR detection of the single-copy Y chromosome marker, *SRY*. HD-derived cells were inferred by subtracting the *SRY* allele count from the total number of human genomes (half the number of the diploid *RPP30* allele count). Male donors were selected based on high *CALR* MAF (mean 50% ±0.2%), indicating pure *CALRmut* grafts, and the confirmed absence of Y chromosome deletion or amplification.

**Figure 5:**
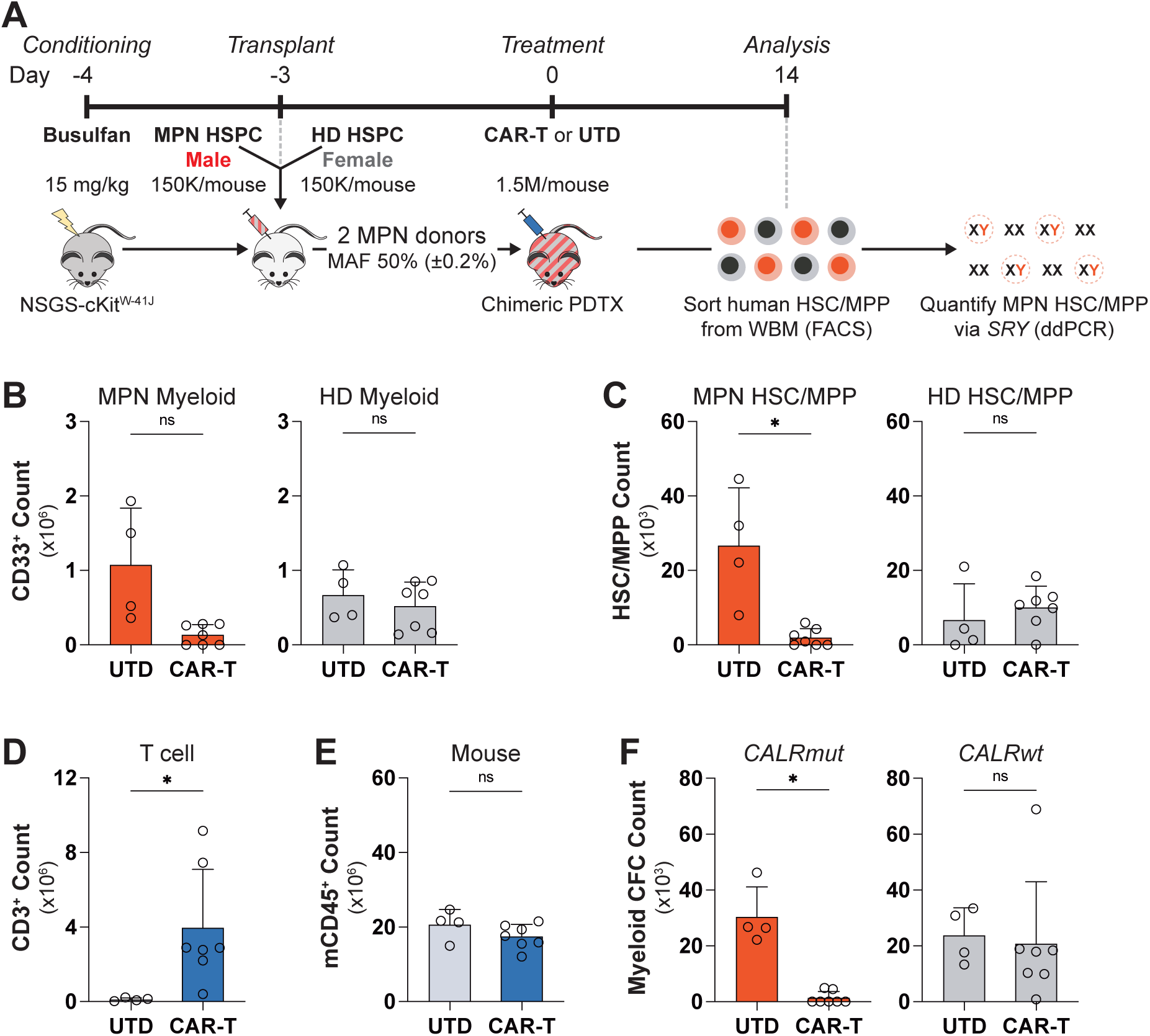
huAB2 CAR-T cells selectively eliminate MPN hematopoiesis while preserving normal hematopoiesis *in vivo*. **A)** Experimental design for *in vivo* evaluation of huAB2 CAR-T cell selectivity using chimeric PDTX mice generated with mixed donor grafts containing both MPN HSPCs and HD HSPCs. Male MPN donors and female HD were used to enable tracking of MPN hematopoiesis via *SRY* genotyping. **B)** Mean human CD33^+^ myeloid cell counts in bone marrow of chimeric PDTX mice 14 days after treatment with CAR-T cells (n = 7) or UTD (n = 4), stratified by donor origin (male/CALRmut/MPN versus female/CALRwt/HD). **C)** Mean human HSC/MPP counts in post-treatment chimeric PDTX bone marrow, stratified by donor origin. **D)** Mean human T cell counts in post-treatment chimeric PDTX bone marrow. **E)** Mean recipient-derived murine leukocyte counts in post-treatment chimeric PDTX bone marrow. **F)** Myeloid CFC assays from post-treatment chimeric PDTX bone marrow. Mean CFC counts are shown for CAR-T- and UTD-treated mice, stratified by *CALRmut* and *CALRwt* genotypes. Error bars represent SD and each point represents an individual mouse. * p < 0.05, ** p < 0.01, *** p < 0.001, ns = not significant (p ≥ 0.05) (t-test).

In the bone marrow of huAB2 CAR-T-treated chimeric PDTX mice, male MPN donor-derived HSC/MPPs were nearly eliminated, with a corresponding reduction in MPN-derived CD33^+^ myeloid cells compared to UTD-treated controls (Fig. 5B,C). In contrast, female HD-derived hematopoietic cells were preserved at levels comparable to those in UTD-treated controls. Despite robust CAR-T cell expansion in recipient mice (Fig. 5D), recipient-derived murine leukocyte numbers were unaffected, indicating the absence of non-specific, off-target toxicity (Fig. 5E). Selective depletion of male MPN hematopoiesis with preservation of female HD progenitor cell function was confirmed by CFC assays and sex genotyping of individual colonies. Following huAB2 CAR-T treatment, residual CFCs were almost exclusively of female HD origin, whereas colonies from UTD-treated controls were derived equally from male MPN and female HD cells concordant with the ratio in the original grafts (Fig 5F). Therefore, huAB2 CAR-T cells readily eliminate *CALRmut* MPN HSC/MPPs *in vivo* while preserving coexisting normal hematopoietic elements.

### huAB2 CAR-T cells eliminate functional disease-initiating MPN HSCs *in vivo*

To determine whether huAB2 CAR-T treatment eliminates functional MPN HSCs capable of long-term disease reconstitution, we performed secondary transplantation assays using whole bone marrow harvested from primary MPN PDTX mice 14 days after treatment with huAB2 CAR-T cells or UTD controls (as shown in Fig. 4). Bone marrow harvested and pooled from both femurs of male MPN donor HSPC-engrafted recipients was mixed with 50,000 female HD HSPCs (added as an internal control for human engraftment) and transplanted into NSGS-cKit^W-41J^ secondary recipients (Fig. 6A). Human hematopoietic engraftment was serially monitored in peripheral blood over 8 weeks post-transplant. In all cohorts, co-transplanted HD-derived HSPCs successfully engrafted secondary recipients, confirming the technical success of transplantation (Fig. 6B). The secondary recipient of bone marrow from a UTD-treated primary PDTX mouse showed persistent MPN reconstitution, with 25% of human hematopoiesis derived from male *CALRmut* clones. In contrast, no *CALRmut*-derived hematopoiesis was detected in the secondary recipient of bone marrow from a huAB2 CAR-T-treated primary PDTX mouse (Fig. 6C). These data indicate complete loss of *CALRmut* reconstituting activity following huAB2 CAR-T therapy. Together, these results show that huAB2 CAR-T cells eliminate not only immunophenotypic MPN HSC/MPP populations but also the functional MPN reconstituting units responsible for long-term disease propagation, providing *in vivo* evidence for eradication of disease-propagating MPN HSCs.

**Figure 6:**
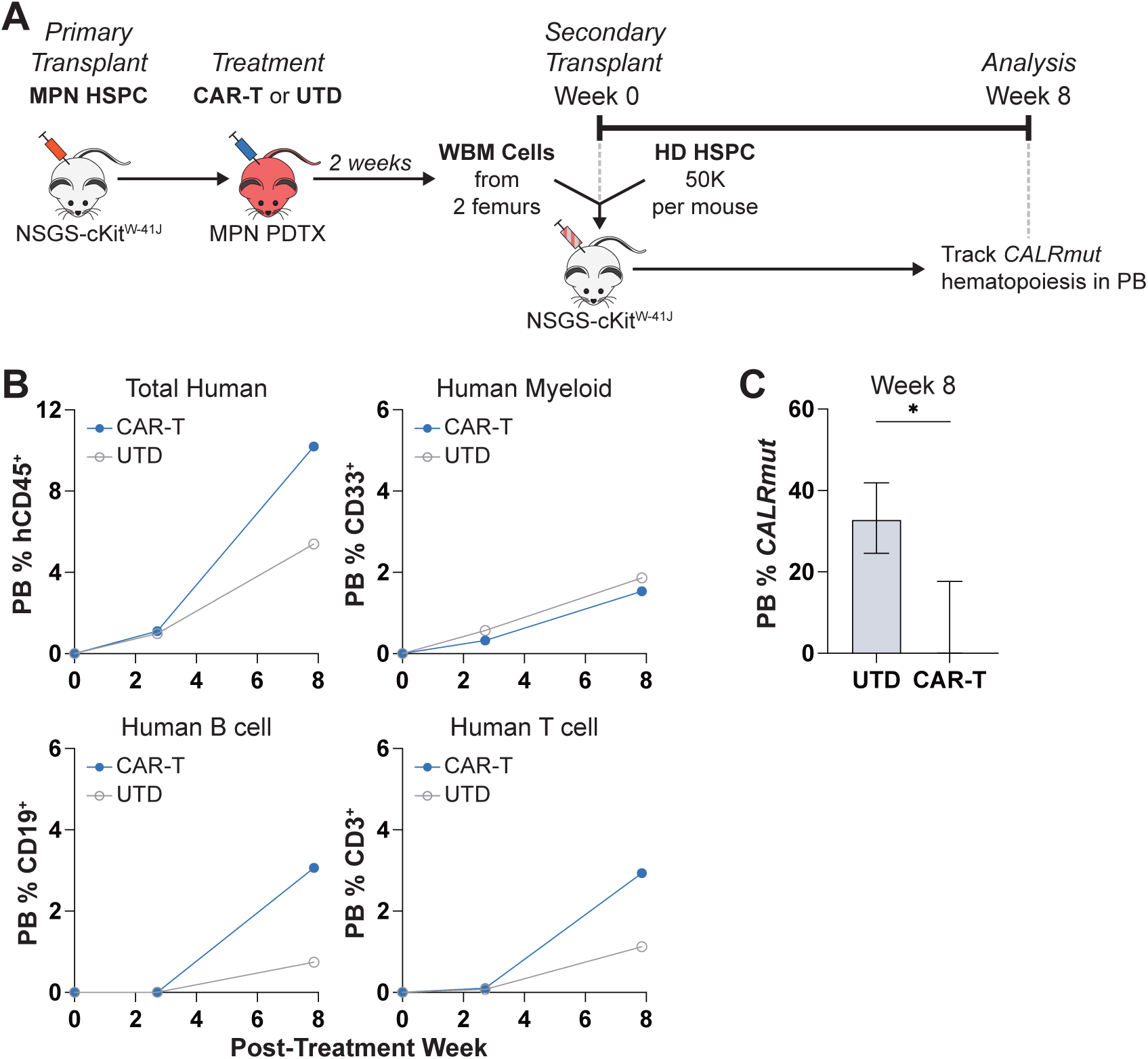
huAB2 CAR-T cells eliminate functional MPN HSCs *in vivo*. **A)** Experimental design for functional assessment of MPN HSCs via secondary transplant of whole bone marrow (WBM) cells from CAR-T- or UTD-treated primary MPN PDTX donors (from Figure 4) into immune-deficient recipient mice. Each secondary graft consisted of WBM cells isolated from two femurs of a primary MPN PDTX mouse mixed with HD HSPC as an internal control for overall engraftment capacity. **B)** Serial assessment of total human CD45^+^ leukocyte, CD33^+^ myeloid, CD19^+^ B cell, and CD3^+^ T cell engraftment by flow cytometry in secondary recipients of CAR-T- or UTD-treated MPN PDTX WBM. **C)** Detection of *CALRmut* alleles by ddPCR, expressed as a percentage of total human genomes, in secondary recipients of CAR-T- or UTD-treated MPN PDTX bone marrow. Error bars represent 95% CI. * p < 0.05 (Poisson exact test).

## Discussion

Although MPNs are sustained by mutated HSCs, existing therapies rarely eliminate this disease-propagating compartment, leaving allogeneic HSC transplantation as the only established curative option (36). Here, we show that recurrent *CALR* frameshift mutations create a uniquely actionable cell-surface neoantigen (CALRmut) that enables selective immunologic elimination of MPN HSCs *in vivo*. Using huAB2, a humanized CALRmut-specific monoclonal antibody, as the targeting domain for CAR-T cells, we show efficient recognition of CALRmut in its physiologic MPL-associated context on MPN HSCs, robust *in vivo* cytotoxicity against primary patient-derived MPN hematopoiesis, selective preservation of coexisting normal hematopoiesis, and complete elimination of functional MPN reconstituting cells as assessed by secondary transplantation. Collectively, these findings establish CALRmut as an accessible and biologically constrained therapeutic target on disease-sustaining HSCs and demonstrate the feasibility of HSC-directed immunotherapy as a transplant-alternative strategy for MPN cure.

HSC-directed immunotherapy for *CALRmut* MPNs requires display of the MPN-defining target antigen on the surface of disease-propagating HSCs but not normal HSCs. Prior studies have shown surface CALRmut staining within CD34^+^ HSPC subpopulations, including immunophenotypic HSCs, in mouse models and in patients with *CALRmut* MPNs (37–39), suggesting that this prerequisite might be met. However, the reported shifts in CALRmut fluorescence intensity were modest and largely overlapped with negative controls, suggesting that a substantial fraction of immunophenotypic MPN HSCs lack detectable surface CALRmut. Moreover, these studies did not address whether CALRmut-expressing HSCs encompass the full pool of functional MPN-propagating cells. Consequently, it has remained unclear if a reservoir of MPN HSCs lacking surface CALRmut exists that could evade immunologic targeting and preclude curative therapy.

Using antibody-based labeling and *in vivo* targeting, we demonstrate that CALRmut is broadly and consistently displayed on disease-propagating MPN HSCs in both human samples and physiologic murine models. Functional transplantation assays demonstrating the selective loss of *CALRmut* reconstituting activity following huAB2 CAR-T treatment of MPN PDTX mice argue against the existence of a significant cryptic population of MPN HSCs lacking surface CALRmut. These findings establish surface CALRmut display as a defining and therapeutically exploitable feature of MPN HSCs, satisfying a fundamental requirement for HSC-directed immunotherapy.

The accessibility of MPN HSCs to immunologic targeting *in vivo* has been uncertain. CALRmut requires MPL for cell-surface display and acts as a pseudoligand that drives MPL signaling. However, the role of MPL signaling in normal and malignant HSC biology remains incompletely understood. Although genetic studies have implicated MPL in the regulation of HSC quiescence (40,41), the profound megakaryopoietic defects in these models raise the possibility that the loss of HSC quiescence reflects indirect loss of megakaryocyte-derived niche factors (42–44) rather than HSC-intrinsic MPL signaling. Importantly, if CALRmut-mediated MPL signaling were to enforce HSC quiescence, constitutive activation of this pathway would be predicted to reduce, rather than enhance, *CALRmut* HSC fitness, as clonal dominance requires increased self-renewal and/or survival relative to wild-type HSCs. This conceptual tension has left unresolved whether CALRmut-mediated MPL signaling is functionally engaged in MPN HSCs *in vivo*. Furthermore, it remains unknown if the specialized hematopoietic microenvironments of the bone marrow and spleen permit effective access to MPN HSCs by antibody- or T cell-based immunotherapies.

These uncertainties are directly addressed by data showing that CALRmut is displayed on the surface of MPN HSCs and targetable in hematopoietic microenvironments, enabling immunologic elimination of functional MPN reconstituting cells. The efficacy of huAB2 CAR-T cells *in vivo* indicates that CALRmut-mediated MPL signaling is not merely permissive but actively maintained in MPN HSCs, and that quiescent HSCs are not immunologically invisible when they express a surface antigen that drives malignant fitness. These findings revise prevailing assumptions about HSC immune privilege and indicate that physiologic target antigen density and quiescence may be less limiting for HSC-directed immunotherapy than previously anticipated.

A critical finding is the preservation of normal hematopoiesis seen following huAB2 CAR-T therapy. In chimeric PDTX models containing both *CALRmut* MPN and normal human HSCs, huAB2 CAR-T cells selectively eliminated malignant HSCs while sparing co-existing normal HSCs. Consistent with this selectivity, no evidence of collateral damage to recipient-derived murine hematopoiesis was seen in huAB2 CAR-T cell recipients. Remarkably, even in MPN PDTX models derived from patient samples in which normal HSCs were initially undetectable, *CALRwt* hematopoiesis could emerge following huAB2 CAR-T treatment, demonstrating the potential for selective survival and functional expansion of residual normal HSCs under therapeutic pressure.

Secondary transplantation assays further confirmed that all residual long-term human reconstituting activity after huAB2 CAR-T treatment was exclusively derived from normal HSCs. Therefore, strict CALRmut specificity translates into functional selectivity at the HSC level that fundamentally distinguishes this approach from allogeneic HSCT, where curative efficacy is inseparable from ablation of normal hematopoiesis and prolonged immunosuppression.

This study shows that eradication of MPN at its stem cell root is an achievable therapeutic goal and provides a strong rationale for clinical translation. Other CALRmut-targeted therapeutic strategies are in development (38,45–48), and their effects on disease-propagating MPN HSCs remain to be determined. huAB2 CAR-T cells efficiently accessed and eradicated MPN HSCs *in vivo*, countering longstanding assumptions that HSC quiescence, niche biology, or tissue localization would limit the effectiveness of cellular immunotherapy. The loss of functional *CALRmut* hematopoiesis in CFC and secondary transplantation assays argues against antigen escape at the level of MPN HSCs and supports surface CALRmut as a stable and obligate feature of MPN HSC fitness. These findings establish a foundation for curative non-transplant therapeutic strategies for *CALRmut* MPNs. Ongoing and future first-in-human studies will define the efficacy, durability, and safety of CALRmut-targeted immunotherapeutic effects in patients.

## Methods

### Human samples and cell lines

Peripheral blood specimens from patients with MPNs and healthy controls, as well as cord blood units, were obtained at Weill Cornell Medicine (WCM) using procedures approved by the WCM institutional review board (IRB). Written informed consent was obtained from all participating donors in accordance with the Declaration of Helsinki. Each sample was de-identified at the site of collection by clinical research staff and assigned a unique donor ID. This donor ID was linked to patient age, diagnosis, treatment history, and other clinical data including next-generation sequencing results. De-identified data were accessed through the WCM MPN Center Research Data Repository (RDR) as previously described (49).

Peripheral blood mononuclear cells (PBMCs) were partitioned from donor blood by density-gradient separation using Ficoll-Paque PLUS (Cytiva, Uppsala, Sweden). CD34^+^ HSPC and CD34-depleted fractions were obtained from PBMCs by immunomagnetic separation (Miltenyi Biotec, Bergisch Gladbach, Germany) and cryopreserved. Human T cells were isolated from the CD34-depleted fractions by negative selection using the pan T cell isolation kit (Miltenyi Biotec).

HSC/MPPs were purified from the CD34^+^ fractions by FACS using previously described methods (50) Ba/F3 cells (DSMZ, Braunschweig, Germany) double-expressing human MPL with either human mutant CALRdel52 (Ba/F3mut) or wild-type CALR (Ba/F3wt) were generated by lentiviral transduction and double antibiotic selection. Cells were maintained in RPMI-1640 (Thermo Fisher Scientific, Waltham, MA, USA) supplemented with 10% fetal bovine serum (FBS) (Corning, Corning, NY, USA) and 10 ng/ml recombinant mouse IL3 (Preprotech, Rocky Hill, NJ, USA).

### Antibody discovery

CALRmut-specific monoclonal antibodies were generated from wild-type mice (Curia, Albany, NY, USA) by immunization with synthetic CALRmut neopeptides conjugated to carrier proteins. Hybridomas were screened for selective binding to full-length CALRmut protein and CALRmut neopeptide by ELISA and to Ba/F3mut cells by FACS. Positive clones were counter-screened to eliminate non-specific binders via FACS using Ba/F3wt and Ba/F3 parental cells. Lead candidates were evaluated by ELISA for binding to heparin and baculovirus particles to eliminate any that exhibited polyreactivity, then fully sequenced. Antibody binding to biotinylated CALRmut neopeptide captured on streptavidin biosensors was assessed with antibody dilution series by biolayer interferometry (BLI) using the Octet RED96e system (Sartorius, Göttingen, Germany). Lead candidates were affinity ranked to select the highest-affinity CALRmut-specific antibody (AB2), which was humanized by grafting its murine complementarity-determining regions onto human antibody variable region germline sequences.

### CAR-T cell production

A second-generation human CAR was engineered using the scFv of humanized AB2 (huAB2) tagged by the HA epitope and fused to the human CD8α hinge/transmembrane domains, the 4-1BB costimulatory domain, and the intracellular human CD3ζ signaling domain. Human T cells from peripheral or cord blood were activated overnight using CD3/CD28 microbeads (Miltenyi Biotec) and transduced with CAR-containing lentiviral supernatants to generate huAB2 CAR-T cells. After microbeads were removed on day 3 post-transduction, CAR-T cells or matched untransduced T cell controls (UTD) were expanded *ex vivo* in Yssel’s T cell medium (GeminiBio, Sacramento, CA, USA), harvested on day 12 post-transduction, then cryopreserved.

### Mice

All mice were maintained in the WCM Animal Facility according to protocols approved by the WCM Institutional Animal Care and Use Committee. Mice were allocated to age- and sex-matched cohorts then randomly assigned to treatment. Investigators were blinded to treatment allocation during data collection and analysis.

### Murine syngeneic *CALRmut* MPN model

Heterozygous floxed *CALRdel52* mice (*CALR^+/FL^*) were crossed with *vav*Cre transgenic mice (Jackson Laboratories, Bar Harbor, ME, USA) to generate *vav*Cre-*CALR^+/MUT^* mice. Fluorescent conjugates of AB2 and murine IgG2a-kappa isotype control (Biolegend) were obtained using Alexa Fluor 647 succinimidyl ester (Thermo Fisher Scientific). Fluorescently-labeled AB2 or isotype control at 5mg/kg in PBS was infused into *CALR^+/FL^*or *CALR^+/MUT^* mice (aged 14 to 18 weeks) via retro-orbital injection. Blood for complete blood counts (CBC) and flow cytometry analysis was obtained by retro-orbital bleeding. Mice were euthanized 48 hours after infusion. Lineage-negative (Lin^-^) cells were isolated from harvested whole bone marrow via negative selection using the mouse lineage depletion kit (Miltenyi Biotec) for further FACS and immunofluorescence analysis.

### Murine CALRmut tumor xenograft model

NSGS mice (aged 6 weeks) were implanted via tail vein injection with 5,000 Ba/F3mut cells per mouse in PBS with 195,000 syngeneic NSGS bone marrow cells added as a carrier. One day after Ba/F3mut implantation, mice were treated with 750,000 huAB2 CAR-T cells or UTD-T cells via tail vein injection. Mice were euthanized on post-treatment day 14 and peripheral blood, whole bone marrow and spleen harvested to enumerate mCD45^+^MPL^+^ Ba/F3mut tumor, hCD45^+^ human T cell, and mCD45^+^MPL^-^ host subpopulations. Murine and human CD45 are designated mCD45 and hCD45, respectively.

### Human patient-derived tumor xenograft models

NSGS-cKit^W-41J^ mice (generated by backcrossing NSGS mice into NBSGW, aged 6 weeks) were conditioned with one 15mg/kg dose of busulfan. The following day, previously cryopreserved HSPCs from MPN donors were implanted via tail vein injection either alone (to generate MPN PDTX mice) or in a 1:1 mixture with HSPCs from HD (to generate chimeric PDTX mice). Each MPN HSPC sample was divided equally to implant at least 3-4 mice to split between treatment groups. Each mouse received a total of 300,000 HSPCs in PBS. Three days after HSPC implantation, mice were treated with 1.5 million huAB2 CAR-T cells or UTD-T cells via tail vein injection. Mice were euthanized on post-treatment day 14 and whole bone marrow and spleen harvested to enumerate human hematopoietic subpopulations.

To assess post-treatment functional hematopoietic reconstituting activity, whole bone marrow obtained from crushing two femurs of each primary treated MPN PDTX mouse was mixed with 50,000 HSPCs from HD and implanted via tail vein injection into secondary busulfan-conditioned NSGS-cKit^W-41J^ recipients. Each primary MPN PDTX bone marrow sample was implanted into one secondary recipient. Serial blood samples for flow cytometry and ddPCR analysis were obtained from secondary recipients by retro-orbital bleeding.

### Cytotoxicity assays

HSPCs isolated from MPN or HD peripheral blood samples were plated at 1 million cells/ml in StemSpan SFEM (StemCell Technologies, Vancouver, Canada) supplemented with 25ng/ml each of recombinant human TPO, SCF, and FLT3-ligand (Preprotech). One day later, cells were treated with huAB2 CAR-T cells or UTD-T cells at a 1:1 effector-to-target ratio. At 72 hours post-treatment, CD3^-^ target cell counts were obtained by FACS from each culture and normalized to the respective counts obtained from untreated control cultures to obtain target cell fold changes.

### Myeloid colony forming cell assays

Bone marrow cells from treated MPN or chimeric PDTX mice were plated in duplicate in MethoCult SF H4436 methylcellulose media (StemCell Technologies) at 50,000 cells per 35mm dish. After 12 days, all colonies per culture were counted and classified using an inverted brightfield microscope (Zeiss, Oberkochen, Germany). Up to 20 individual colonies were picked from each culture for *CALR* genotyping as described below.

### Flow cytometry

Antibodies (with clones in parentheses) for the following human antigens were obtained from Biolegend (San Diego, CA, USA): CD3 (HIT3a), CD34 (561), CD38 (HIT2), CD45 (HI30), CD45RA (HI100), CD90 (5E10), granzyme B (GB11), MPL (S16017A), perforin-1 (δG9). Antibodies for the following mouse antigens were obtained from Biolegend: B220 (RA3-6B2), CD3 (17A2), CD16/32 (93), CD41 (MWReg30), CD45 (30-F11), CD48 (HM48-1), CD150 (TC15-12F12.2), cKit (2B8), Sca1 (D7), Ter-119 (TER-119). Anti-human CD33 (P67.6) and CD4 (RPA-T4) antibodies were obtained from BD Biosciences (San Jose, CA, USA). Anti-human CD8 (SFCI21Thy2D3) and CD19 (J3-119) antibodies were obtained from Beckman Coulter Life Sciences (Brea, CA, USA). Anti-human blocking was completed using Human TruStain FcX (Biolegend).

Cells were stained for flow cytometry for 30 minutes in FACS buffer (0.2% BSA and 2mM EDTA in PBS) at 4°C. Viability was assessed by staining live cells with DAPI (Biolegend) prior to analysis, or by staining cells with the Zombie UV fixable viability kit (Biolegend) prior to fixation. For surface CALR staining of primary cells, cells were stained first with fluorescent AB2 for 12 hours in IMDM (Thermo Fisher Scientific) at 37°C, washed twice with FACS buffer, then stained for other surface markers in FACS buffer at 4°C as above. For intracellular perforin-1 and granzyme B staining, cells were fixed and permeabilized using the IntraPrep Permeabilization Reagent (Beckman Coulter) according to the manufacturer’s instructions, then stained for 15 minutes at 21°C. Cell sorting was completed to a mean purity of 98% using a FACSymphony S6 cell sorter (BD Biosciences). Flow cytometry data were visualized and analyzed using FCS Express (De Novo Software, Pasadena, CA, USA). Raw median fluorescence intensity (MFI) values for AB2 staining were adjusted by subtracting MFI of matched samples stained with murine isotype control to obtain background-subtracted AB2 MFI values that are reported in the figures.

### *CALR* MAF and *SRY* copy number quantification

*CALRmut* (including Type 1 *CALRdel52* and Type 2 *CALRins5* mutations), *CALRwt*, *SRY*, and *RPP30* alleles were detected in genomic DNA (gDNA) isolated from sorted cell subpopulations using custom or commercial primer/probe assays (specified below) for ddPCR according to previously described protocols.(50) *CALR* MAF, *SRY* copy number and 95% confidence intervals were estimated using Poisson statistics in R using the rateratio.test package.

Fold changes in absolute numbers of *CALRmut* and *CALRwt* alleles during *in vitro* cytotoxicity assays (Supplementary Fig. S4B) were calculated as follows:

*CALRmut* allele fold change = N_F_ * MAF_F_ / N_0_ * MAF_0_,
*CALRwt* allele fold change = N_F_ * (1 – MAF_F_) / N_0_ * (1 – MAF_0_),

where N_F_ = final cell count, MAF_F_ = final MAF, N_0_ = initial cell count, MAF_0_ = starting MAF.

Male MPN donor-derived human cells were distinguished from female HD-derived human cells in chimeric PDTX (Fig. 5) via *SRY* copy number as follows:

MPN donor-derived cell count = N_T_ * N_SRY_ / N_G_,
HD-derived cell count = N_T_ * (N_G_ – N_SRY_) / N_G_,

where N_T_ = total cell count, N_SRY_ = *SRY* copy number, N_G_ = total number of diploid human genomes (half the copy number of diploid *RPP30*).

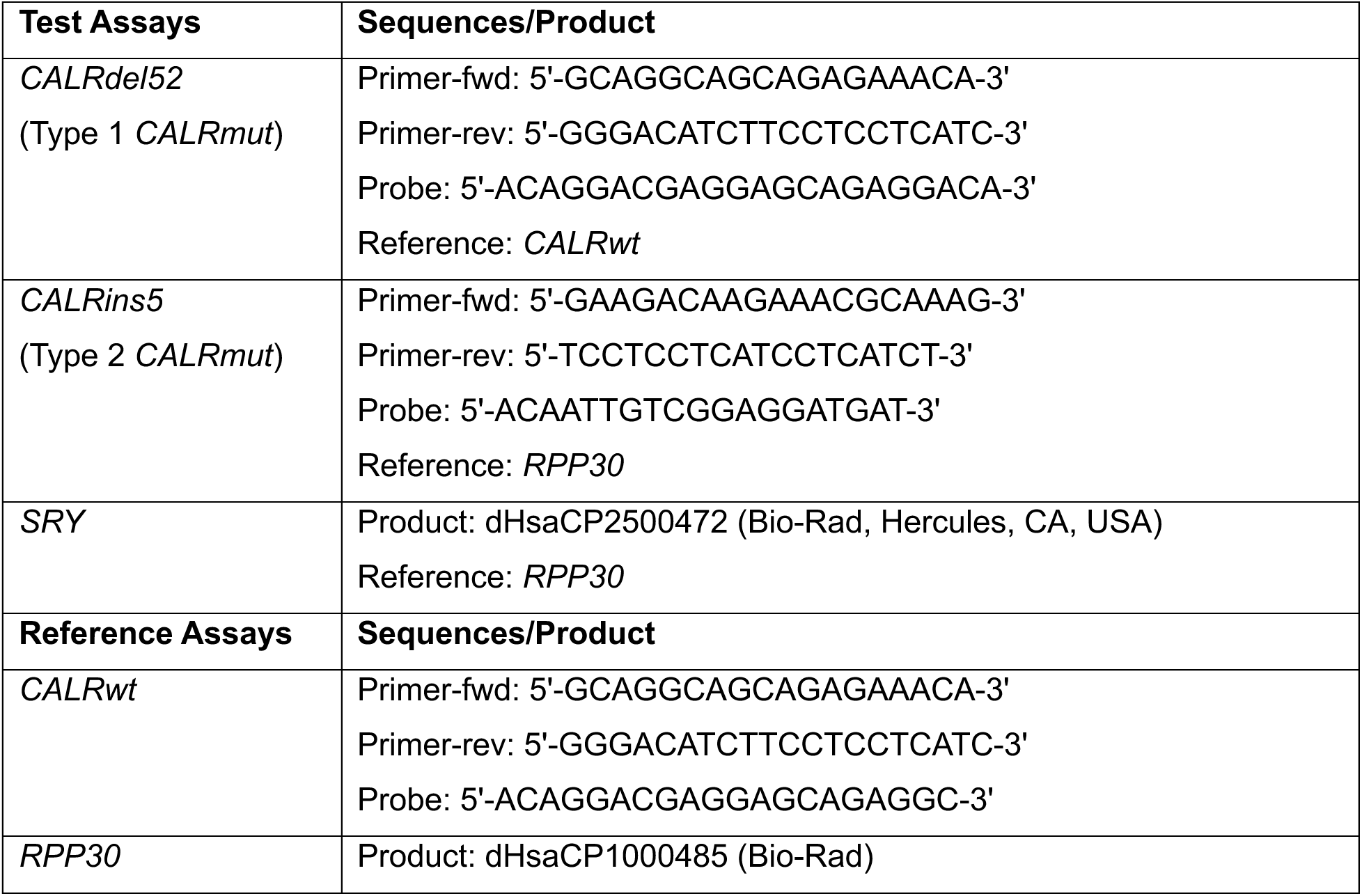

### Histopathological analyses

Immunohistochemistry was completed on 4μm formalin-fixed, paraffin-embedded sections of tissues collected from euthanized mice. Slides were deparaffinized, rehydrated, and subjected to antigen retrieval on a BOND-III automated stainer (Leica Biosystems, Nussloch, Germany). Sections were stained with AB2, counterstained with hematoxylin, and visualized using the BOND Polymer Refine Detection kit (Leica Biosystems) on an inverted brightfield microscope (Zeiss).

Immunofluorescence was completed on murine Lin^-^ cells deposited onto glass coverslips. Cells were fixed with 1% paraformaldehyde, permeabilized with 0.1% Triton-X, stained with fluorescent anti-CD48 and anti-CD150 antibodies, and counter-stained with DAPI. Confocal images were acquired using an inverted Ti2-E microscope (Nikon Instruments, Tokyo, Japan) equipped with a CSU-W1 SoRa module (Yokogawa, Tokyo, Japan) and an ORCA-Fusion BT sCMOS camera (Hamamatsu, Hamamatsu City, Japan). Imaging was performed using a 60x Plan Apo Lambda oil-immersion objective (numerical aperture 1.42). Z-stack images were collected for each field of view and extended depth-of-field (EDF) projections were generated using NIS-Elements software (Nikon Instruments). Image analysis was completed using QuPath.

### Statistical analyses

Dose-response data from BLI and FACS were fit to a 1:1 Langmuir binding model to compute rate constants (k_on_ and k_off_) and equilibrium dissociation constants (K_D_). Statistical comparisons between experimental groups were completed using unequal variances t-test to compare means between unpaired groups, and one-way analysis of variance (ANOVA) with Tukey Honest Significant Difference (HSD) test to compare means across multiple groups. Overall survival of NSGS-Ba/F3mut treatment cohorts was estimated using Kaplan-Meier methods and compared between groups using log-rank tests. All analyzable data points were included. All p-values were derived from two-tailed statistical analyses using an alpha level of 0.05. Except where otherwise specified, all means are plotted with standard deviations. Significance is denoted with asterisks (* p < 0.05, ** p < 0.01, *** p < 0.001). Non-significant comparisons (p ≥ 0.05) are either not labeled or designated “ns.” Nonlinear regression modeling, statistical comparisons, and survival analyses were completed using R and the referenced packages.

## Data availability

The data generated in this study are available upon request from the corresponding author.

## R package references

− R Core Team (2025). _R: A Language and Environment for Statistical Computing_. R Foundation for Statistical Computing, Vienna, Austria. https://www.R-project.org/.
− Baty F, Ritz C, Charles S, Brutsche M, Flandrois JP, Delignette-Muller ML (2015). A Toolbox for Nonlinear Regression in R: The Package nlstools. Journal of Statistical Software, 66(5), 1-21. doi:10.18637/jss.v066.i05
− Dawson C (2025). _ggprism: A ‘ggplot2’ Extension Inspired by ‘GraphPad Prism’_. R package version 1.0.7, https://CRAN.R-project.org/package=ggprism.
− Fay M (2022). rateratio.test: Exact Rate Ratio Test. R package version 1.1, https://CRAN.R-project.org/package=rateratio.test.
− Kassambara A, Kosinski M, Biecek P (2025). _survminer: Drawing Survival Curves using ‘ggplot2’_. R package version 0.5.1, https://CRAN.R-project.org/package=survminer.
− Therneau T (2024). _A Package for Survival Analysis in R_. R package version 3.8-3, https://CRAN.R-project.org/package=survival.
− Wickham H, Averick M, Bryan J, Chang W, McGowan LD, François R, Grolemund G, Hayes A, Henry L, Hester J, Kuhn M, Pedersen TL, Miller E, Bache SM, Müller K, Ooms J, Robinson D, Seidel DP, Spinu V, Takahashi K, Vaughan D, Wilke C, Woo K, Yutani H (2019). Welcome to the tidyverse. Journal of Open Source Software, 4(43), 1686. doi:10.21105/joss.01686

## Acknowledgements

We are grateful to the patients at the Richard T Silver MPN Center for contributing research specimens. We acknowledge Ghaith Abu-Zeinah, Ellen Ritchie, Katie Erdos and Neville Lee for consenting patients and collecting research specimens. We thank Luca Cappelli and Alexandra Satty their contributions to antibody screening and CAR construct development. This research was supported by the Burroughs Wellcome Weill Cornell Physician Scientist Program and the Weill Cornell Department of Medicine Fund for the Future Award (DCC); the American Society of Hematology (ASH) Inclusion Pathway (FCT); the Daedalus Fund for Innovation (GI and JMS); the Cancer Research & Treatment Fund (CR&T) and the National Heart, Lung, and Blood Institute (NHLBI) grant R01 HL175556 (JMS); and the National Center for Advancing Translational Sciences (NCATS) grant UL1 TR002384 to the Clinical and Translational Science Center (CTSC) of the Weill Medical College of Cornell University.

## Contributions

These authors contributed equally: Daniel C Choi, Giovanni Medico.

These authors jointly supervised this work: Giorgio Inghirami, Joseph M Scandura.

DCC designed the study, performed experiments, collected and analyzed data, and wrote the manuscript; GM designed and performed experiments and collected and analyzed data; IVL and EKN performed antibody screening and engineering; CK, FCT, PK, NM, OV, AT, MMY, and UDC performed experiments; PB and MB directed antibody screening and engineering; GI co-conceived the study and designed experiments; JMS co-conceived the study, designed experiments, examined and consented patients, provided research specimens, and wrote the manuscript.

