## Supplementary material for "CAR-T Targeting of Mutant Calreticulin Establishes a Potentially Curative Stem Cell-Directed Therapy for Myeloproliferative Neoplasms": Data Supplement

### Supplementary Data

**Supplementary Table S1: Clinical and demographic characteristics of patient samples used in this study.** Donor samples used for *in vitro* surface CALRmut staining (Fig. 1B,C) and cytotoxic assays (Supplementary Fig. S4A,B) are designated under "*in vitro*." Donor samples implanted to generate MPN PDTX mice (Fig. 4) are designated under "MPN PDTX." Donor samples used in chimeric grafts to generate chimeric PDTX mice (Fig. 5) are designated under "Chimeric PDTX." Abbreviations: ET (essential thrombocythemia), PET-MF (post-ET myelofibrosis), PMF (primary myelofibrosis), Type 1 *CALR* mutation (*CALRdel52*), Type 2 *CALR* mutation (*CALRins5*), ANA (anagrelide), HU (hydroxyurea), IFN (interferon-α), JAKi (JAK2 inhibitor).

Supplementary Table S1

| Donor | Age | Sex | Diagnosis | *CALR* Mutation | *CALRmut* Class | HSPC *CALR* MAF | Co-Occurrent Mutations | Prior Therapies | Active Therapy | *in vitro* | MPN PDTX | Chimeric PDTX |
|---|---|---|---|---|---|---|---|---|---|---|---|---|
| MPD014 | 74 | F | PET-MF | L367fs*46 | Type 1 | 51% | IDH2 | HU, IFN, trial | JAKi | X | | |
| MPD034 | 77 | M | PMF | L367fs*46 | Type 1 | 50% | ASXL1, SF3B1 | IFN | IFN | | | X |
| MPD037 | 80 | M | PMF | L367fs*46 | Type 1 | 50% | ASXL1, EZH2 | ANA, IFN | JAKi | X | X | |
| MPD092 | 73 | M | PMF | L367fs*46 | Type 1 | 52% | None | JAKi, trial | None | X | X | |
| MPD180 | 74 | F | PET-MF | L367fs*46 | Type 1 | 51% | ASXL1 | JAKi | JAKi | X | X | |
| MPD500 | 67 | F | PET-MF | L367fs*46 | Type 1 | 43% | None | HU, ANA, IFN | JAKi | X | | |
| MPD688 | 67 | F | ET | K385fs*47 | Type 2 | 17% | None | HU, IFN, trial | JAKi | X | | |
| MPD693 | 63 | M | ET | E364fs*49 | Type 1-like | 47% | TET2 | None | None | X | | |
| MPD763 | 70 | M | PMF | L367fs*48 | Type 1-like | 43% | ASXL1, TET2 | HU, JAKi, trial | JAKi | | X | X |

Supplementary Figure S1

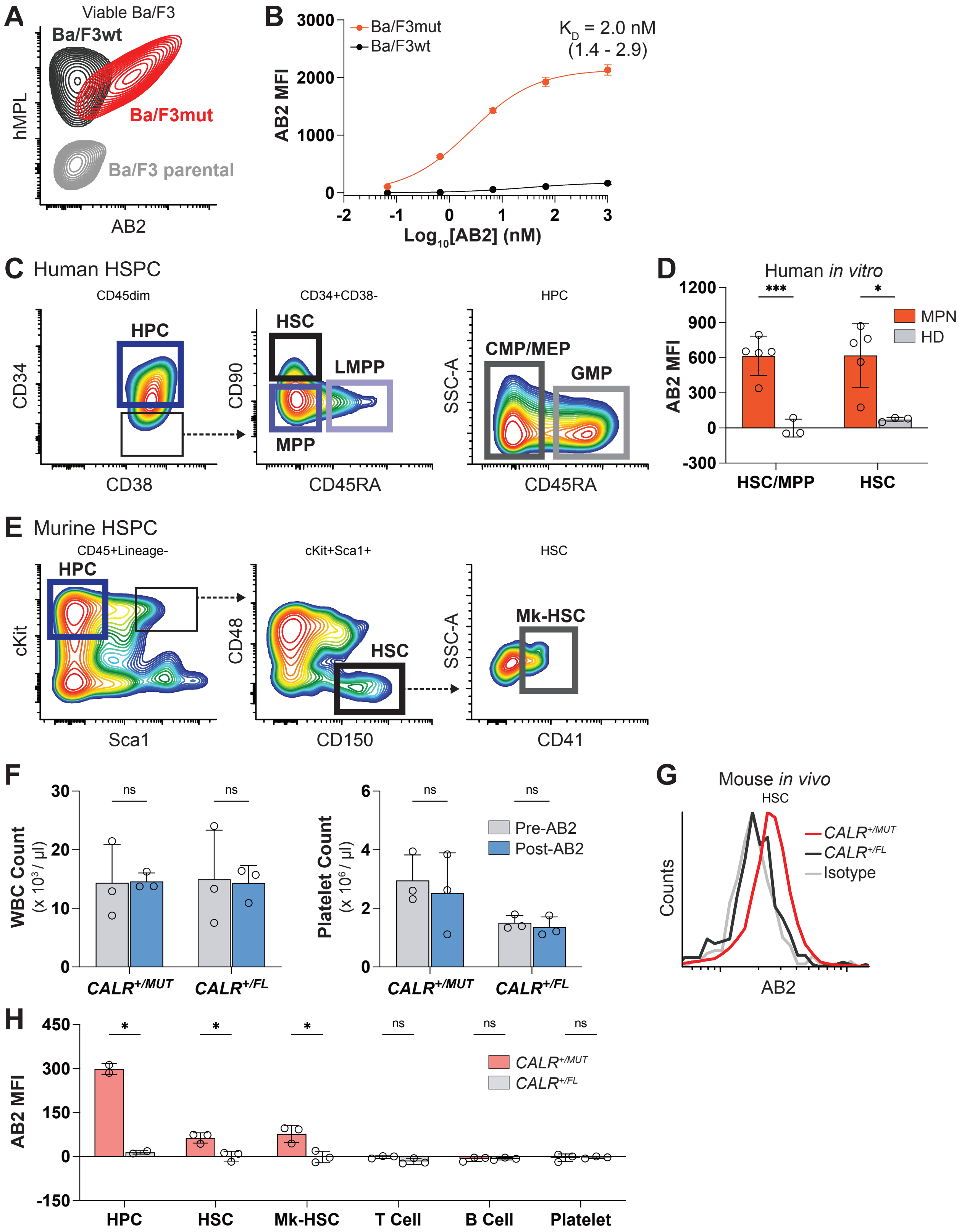

**Supplementary Figure S1: Characterization of CALRmut-expressing cells for evaluation of AB2 targeting. A)** Representative FACS plots showing surface expression of human MPL and CALR (detected by AB2 immunostaining) on Ba/F3mut, Ba/F3wt, and Ba/F3 parental cells. **B)** Nonlinear regression analysis of AB2 specific binding to Ba/F3mut or Ba/F3wt cells. **C)** Flow cytometry gating strategy for identification of human hematopoietic progenitor cells (HPC), hematopoietic stem cells (HSC), multipotent progenitor cells (MPP), lymphoid-primed multipotent progenitor cells (LMPP), granulocytic/monocytic progenitor cells (GMP), and common myeloid, megakaryocytic, and erythroid progenitor cells (CMP/MEP) within $CD34^{+}$ HSPCs from patients with MPNs. **D)** Background-subtracted MFI of *in vitro* AB2 staining of immunophenotypic HSCs within the HSC/MPP compartment from MPN donors or HD. **E)** Gating strategy for identification of murine HPC, HSC, and megakaryocyte-biased HSC (Mk-HSC) within lineage-negative cells from bone marrow of $CALR^{+/MUT}$ mice. **F)** Mean leukocyte counts (WBC, left) and platelet counts (right) in $CALR^{+/MUT}$ or $CALR^{+/FL}$ mice measured prior to (gray) or 48 hours post (blue) *in vivo* AB2 infusion. **G)** Representative histogram showing *in vivo* fluorescent AB2 labeling of bone marrow HSCs in $CALR^{+/MUT}$ (red) or $CALR^{+/FL}$ (black) mice 48 hours after AB2 infusion, relative to background labeling by a murine IgG2a isotype control (gray). **H)** Background-subtracted MFI for *in vivo* AB2 staining of bone marrow stem and progenitor cells, as well as peripheral blood T cells, B cells, and platelets, in $CALR^{+/MUT}$ mice or $CALR^{+/FL}$ control mice at 48 hours after AB2 infusion. Error bars represent SD and each point represents one donor (*in vitro*) or mouse (*in vivo*). * $p < 0.05$, ns = not significant ($p \geq 0.05$) (t-test).

Supplementary Figure S2

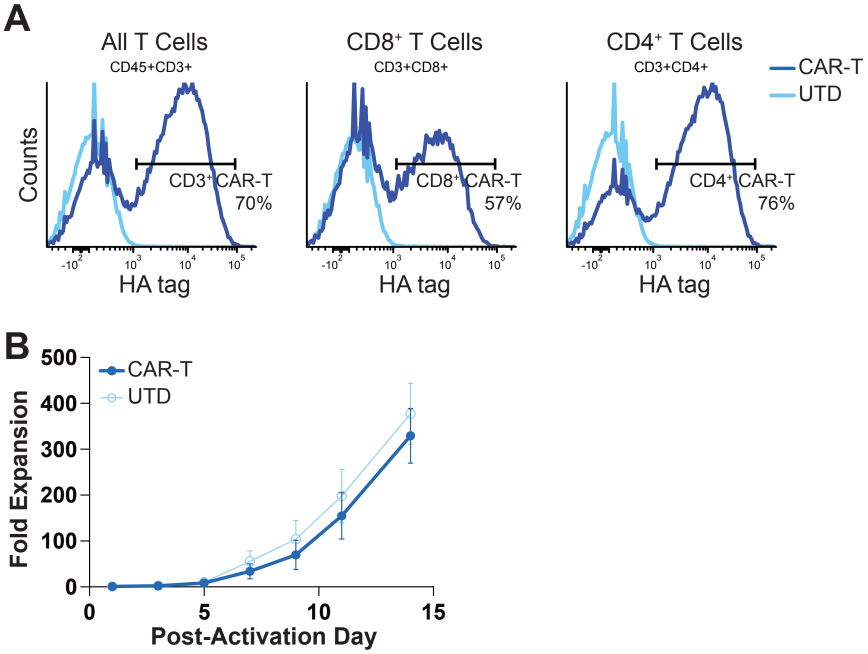

**Supplementary Figure S2: Engineering huAB2 CAR-T cells. A)** Representative flow cytometry histograms showing expression of the HA epitope on huAB2 CAR-T cells (dark blue) or UTD-T cells (light blue). CAR transduction efficiency is shown for the total $CD3^+$ CAR-T population (left) as well as the $CD8^+$ (middle) and $CD4^+$ (right) CAR-T subsets. **B)** *In vitro* expansion kinetics following CD3/CD28 stimulation of huAB2 CAR-T cells (dark blue) and the parent UTD cells (light blue) obtained from 3 distinct donors. Quantified data are presented as means with error bars representing SD.

Supplementary Figure S3

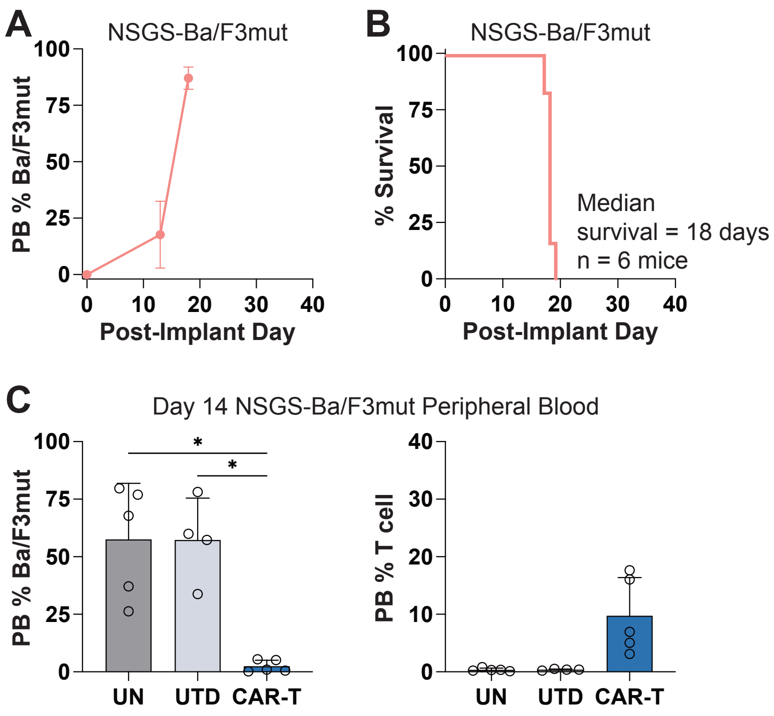

**Supplementary Figure S3: NSGS-Ba/F3mut murine tumor model manifests aggressive and lethal disease. A)** Expansion of mean Ba/F3mut cell counts in peripheral blood (PB) following implantation into NSGS mice. **B)** Kaplan-Meier survival analysis showing rapid mortality of NSGS-Ba/F3mut mice within 3 weeks post-implantation. **C)** Mean PB Ba/F3mut burden (left) and human T cell persistence (right) measured in NSGS-Ba/F3mut mice 14 days after huAB2 CAR-T, UTD, or no (UN) treatment. Error bars represent SD. * $p < 0.05$, non-significant comparisons not shown (t-test).

Supplementary Figure S4

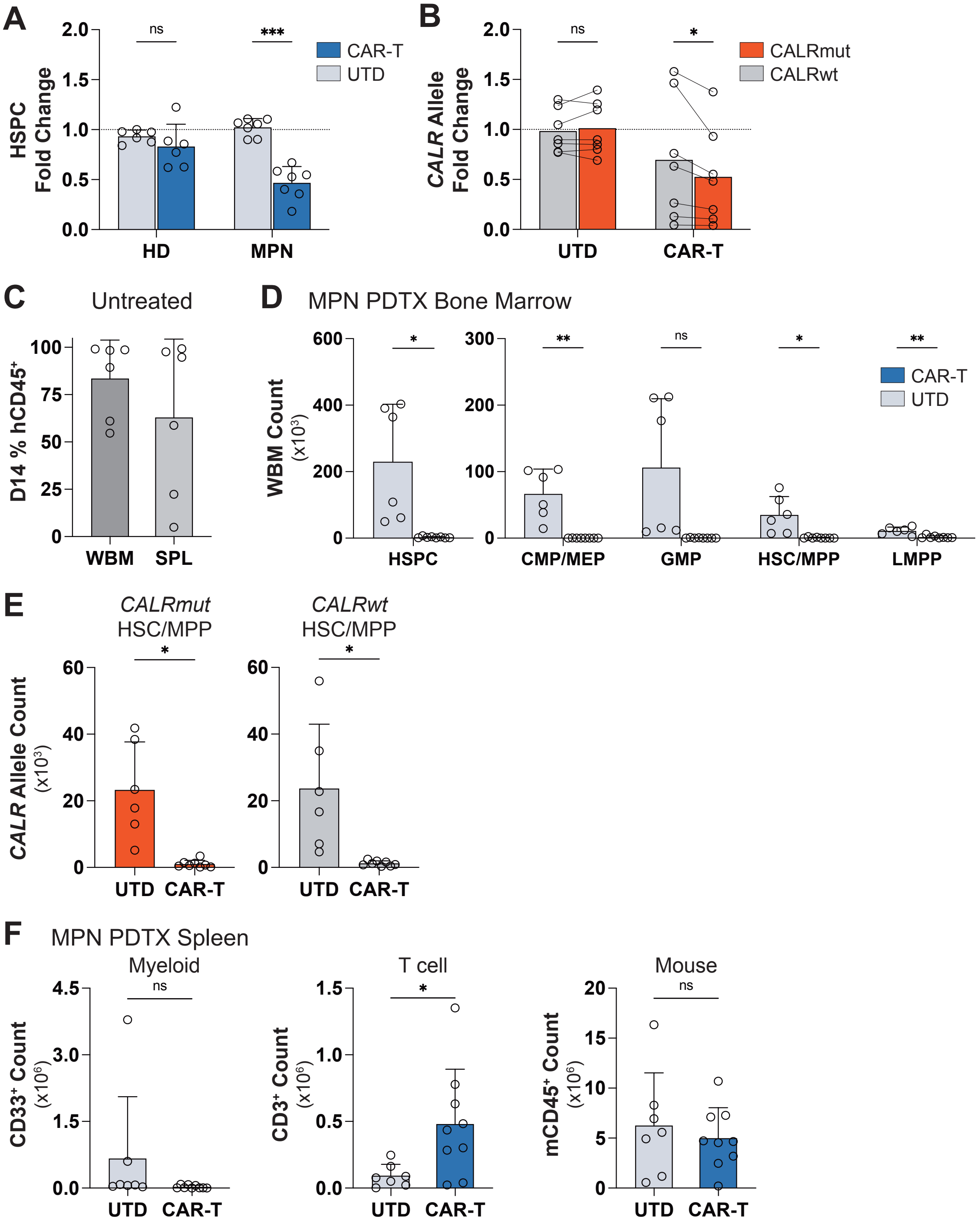

**Supplementary Figure S4: huAB2 CAR-T treatment effects on primary MPN HSPCs. A)** *In vitro* effect of huAB2 CAR-T or UTD cells on mean primary $CD34^+$ HSPC counts from MPN donors or HD, normalized untreated control cultures. **B)** Mean fold change in *CALRmut* (red) or *CALRwt* (gray) allele counts from MPN HSPCs treated with CAR-T or UTD cells relative to untreated controls. Mutant and wild-type *CALR* allele counts from individual MPN donors are paired to highlight the preferential depletion of *CALRmut* alleles following huAB2 CAR-T treatment. **C)** Mean human chimerism (% human $CD45^+$ cells) in the bone marrow (WBM) and spleen (SPL) of MPN PDTX mice 14 days after implantation of MPN HSPCs, without additional treatment. **D)** Total count of $CD34^+$ HSPCs and HSPC subsets in bone marrow of MPN PDTX mice treated with huAB2 CAR-T or UTD cells. **E)** Mean *CALRmut* (left) and *CALRwt* (right) allele counts quantified by ddPCR from HSC/MPPs sorted from MPN PDTX bone marrow after huAB2 CAR-T or UTD treatment. **F)** Mean counts of MPN donor-derived $CD33^+$ myeloid cells, treatment-derived human T cells, and recipient-derived $mCD45^+$ murine leukocytes in the spleens of MPN PDTX mice after huAB2 CAR-T cells or UTD treatment. Error bars represent SD and each point represents one donor (*in vitro*) or mouse (*in vivo*). * $p < 0.05$, ** $p < 0.01$, *** $p < 0.001$, ns = not significant ($p \geq 0.05$) (t-test).
